# Testosterone enhances GLP-1 efficacy at the plasma membrane and endosomes to augment insulin secretion in male pancreatic β cells

**DOI:** 10.1101/2020.05.12.081588

**Authors:** Weiwei Xu, Fiona B. Ashford, Stavroula Bitsi, Lina Schiffer, M.M. Fahd Qadir, Wiebke Arlt, Alejandra Tomas, David J. Hodson, Franck Mauvais-Jarvis

## Abstract

Male mice with elimination of the androgen receptor (AR) in islet β cells (βARKO) exhibit blunted glucose-stimulated insulin secretion (GSIS), leading to hypoinsulinemia and hyperglycemia when challenged with a Western diet. Testosterone activation of an extranuclear AR in β cells potentiates GSIS by amplifying the insulinotropic action of glucagon-like peptide-1 (GLP-1). Here, using a combination of βARKO and β cell-selective GLP-1 receptor knockout mice and their islets, we show that AR activation in β cells amplifies the insulinotropic effect of islet-derived GLP-1. In β cell models expressing cAMP sensors, testosterone enhances the ability of GLP-1, but not that of glucose-dependent insulinotropic polypeptide or glucagon, to produce cAMP. Accordingly, testosterone selectively enhances the ability of GLP-1 to potentiate GSIS. Notably, testosterone enhances GLP-1 production of cAMP at the plasma membrane and endosomes. In male mouse and human islets, the insulinotropic effect of testosterone is abolished following inhibition of the membrane and endosomal cAMP-dependent protein kinase A and exchange protein activated by cAMP islet 2 pathways. Thus, membrane localization of AR enhances the ability of the GLP-1 receptor to produce cAMP, thus increasing glucose-stimulated insulin exocytosis.

**Significance Statement:** This study reveals that testosterone, acting on the androgen receptor (AR) in insulin-producing β cells amplifies the insulinotropic action of glucagon-like peptide-1 (GLP-1) by increasing GLP-1-mediated production of cAMP at the plasma membrane and endosomal compartments, to promote insulin vesicles exocytosis in human β cells. This study establishes a novel biological paradigm in which membrane location of a steroid nuclear receptor enhances the ability of a G protein-coupled receptor to produce cAMP. It has exceptional clinical significance for targeted delivery of testosterone to β cells in the large population of aging and androgen-deficient men who are at increased risk of diabetes.

## Introduction

Early studies showing binding of testosterone to a nuclear protein in prostate led to the establishment of a paradigm in which the androgen receptor (AR) is a nuclear ligand-activated transcription factor (1). We showed that, unlike in classical androgen target tissues, AR is localized outside the nucleus in pancreatic insulin-producing β cells, where it signals in the presence of ligand (2). Activation of the β cell AR by the active metabolite of testosterone, dihydrotestosterone (DHT), enhances glucose-stimulated insulin secretion (GSIS) from cultured male mouse and human islets. We reported that DHT potentiates GSIS by amplifying the insulinotropic action of glucagon-like peptide-1 (GLP-1) in cultured islets (2). Accordingly, when exposed to a western diet, male mice with conditional elimination of the AR in β cells (βARKO) exhibit a defect in GSIS, leading to hypoinsulinemia and hyperglycemia (2). Here, we investigated the mechanism by which DHT acting on AR in β cells enhances GSIS and amplifies the insulinotropic action of GLP-1.

## Results

We generated an inducible β-cell specific ARKO mouse (βARKO^MIP^) by crossing AR^lox^ mice with MIP-CreERT transgenic mice that lacks Cre activity in the hypothalamus (3). The MIP-CreERT transgenic mouse exhibits transgene-driven expression of human growth hormone leading to decreased glucagon secretion and improved insulin sensitivity (4). Therefore, we used AR^lox^ MIP-CreERT mice (without Tam injection) as controls of βARKO^MIP^. We previously validated these controls and showed AR^lox^ (with Tam) and AR^lox^ MIP-CreERT (without Tam) displayed similar glucose tolerance (2). When fed a western diet, βARKO^MIP^ mice developed fed hyperglycemia compared to controls (**Fig. 1A and B**), which was associated with fed hypoinsulinemia as demonstrated by decreased insulin/glucose ratio, an index of β cell failure (**Fig. 1C and D**). We previously showed that DHT amplifies the insulinotropic effect of GLP-1 in cultured mouse and human islets (2). To explore the physiological relevance of these findings *in vivo*, we compared the effect of IP-injected glucose (to partially bypass gut GLP-1 release) versus orally administered glucose (to stimulated gut GLP-1 release) in male control and βARKO^MIP^ mice. Following an IP glucose challenge, βARKO^MIP^ mice developed impaired GSIS (**Fig. 1E**), accompanied by glucose intolerance (**Fig. 1F)**. The defect was selective to glucose sensing as βARKO^MIP^ showed no alteration in arginine-stimulated insulin secretion (**Fig. 1G**). In contrast to the IP glucose challenge, during an oral glucose challenge, βARKO^MIP^ mice exhibited similar glucose tolerance (**Fig. 1H**) and β cell function (I/G ratio at 30 min) (**Fig. 1I**) than littermate controls. Taken together these data suggests that *in vivo*, loss of β cell AR likely impairs the insulinotropic action of islet-derived GLP-1 (reflected by the impaired IP-GTT), but doesn’t seem to alter gut-derived GLP-1 insulinotropic action (reflected by the normal oral glucose challenge).

**Figure 1.**
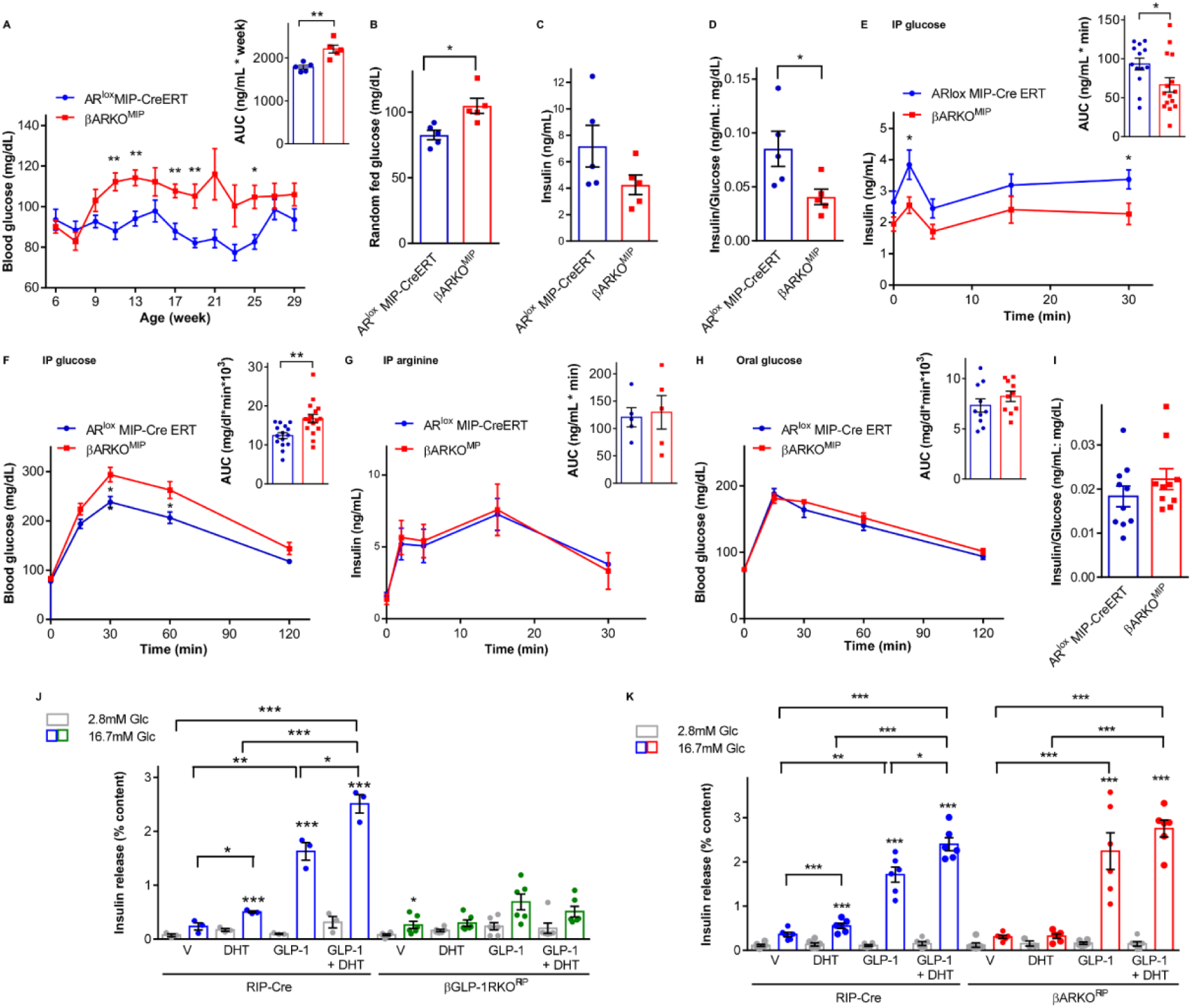
The β cell AR amplifies the insulinotropic effect of islet-derived GLP-1, via β cell GLP-1R. **(A-D)** Data are from βARKO^MIP^ and AR^lox^ MIP-CreERT (control) mice fed a western diet since weaning. **(A)** Random fed blood glucose taken around 10-11am and corresponding area under the curve (AUC) from 9 to 29 weeks. **(B)** & **(C)** Random fed blood glucose and insulin measured at 25 weeks. **(D)** Insulin:glucose index of insulin deficiency from **(B)** & **(C). (E)** IP-GSIS (3 g/kg) with corresponding insulin AUC. **(F)** IP-GTT (2 g/kg) with corresponding glucose AUC. **(G)** IP-ASIS (1 g/kg) with corresponding insulin AUC. **(H)** Oral-GTT (2 g/kg) with corresponding glucose AUC. **(I)** Insulin:glucose ratio at 30 minutes into the Oral-GTT. Mice were studied at 23-35 weeks of age (n = 10-15). **(J-K)** GSIS measured in static incubation in normal chow fed RIP-Cre (control), **(J)** βGLP-1RKO^RIP^ islets and **(K)** βARKO^RIP^ islets treated with DHT (10nM) and GLP-1 (10nM) for 40 minutes. Values represent the mean ± SE of n= 2 mice/group measured in triplicate. * P < 0.05, ** P < 0.01, *** P < 0.001.

To further explore the interaction between DHT and islet-derived GLP-1, we assessed GSIS in static incubation in islets from male control and β cell-specific GLP-1R knockout (βGLP1RKO^RIP^) mice. We used the RIP-Cre transgenic mouse to generate βGLP1RKO^RIP^ and βARKO^RIP^ for cultured islet studies as they provide more complete recombination than using MIP-CreERT transgenic mice (2). Note that mice expressing the RIP-Cre transgene develop impaired insulin secretion (2, 5) and it is recommended that studies involving crosses between floxed mice and RIP-Cre mice use the RIP-Cre as the control group rather than the littermate floxed mice. Consistent with results observed *in vivo*, at 16.7 mM glucose, control islets exposed to DHT showed increased GSIS compared to those exposed to vehicle (**Fig. 1J**). Further, at 16.7 mM glucose, DHT also amplified the insulinotropic action of exogenous GLP-1 on GSIS compared to vehicle-treated islets (**Fig. 1J**). In contrast, in βGLP1RKO^RIP^ islets, DHT alone had no effect on GSIS, and did not amplify the insulinotropic action of exogenous GLP-1 (**Fig. 1J**). Thus, DHT amplifies the insulinotropic action of islet-derived and exogenous GLP-1, and this effect requires the presence of the GLP-1R in β cells. In a parallel set of experiments, we assessed the consequences of loss of β cell AR on GLP-1 insulinotropic action, using male islets from control and βARKO^RIP^ mice. As expected, at 16.7 mM glucose, DHT alone enhanced GSIS and amplified the insulinotropic action of exogenous GLP-1 in control islets (**Fig. 1K**). In contrast, DHT failed to enhance GSIS alone or in the presence of GLP-1 in βARKO^RIP^ islets (**Fig. 1K**). Note that GLP-1’s ability to enhance GSIS was unaltered in βARKO^RIP^ compared to control islets (**Fig. 1K**).

Thus, in male β cells, DHT action on the AR amplifies the insulinotropic action of islet-derived and exogenous GLP-1 acting on the GLP-1R. However, the β cell AR is not essential to the insulinotropic action of exogenous GLP-1 acting on its receptor.

Testosterone (T) is a pro-hormone that requires local conversion to DHT in target tissue via action of the enzyme 5α-reductase (5α-R) to activate AR (6). The observation that βARKO^MIP^ mice exhibit decreased IP-GSIS (**Fig.1**) suggests that testosterone is converted to DHT in male mouse islets *in vivo*. Indeed, using male human pancreas sections, we previously showed that the 5α-R type1 isoform (SRD5A1) is expressed in male β cells. We showed that cultured male human islets exposed to T produce DHT and accordingly, exposure to 5α-R inhibitors inhibited T conversion into DHT (7). To validate T intracrine activation in the mouse, we treated cultured islets from male wild-type mice with T and quantified the conversion of T to DHT by ultra-high-performance liquid-chromatography tandem mass spectrometry (UHPLC-MS/MS). After treatment of cultured islets with T, we detected DHT in the culture supernatant (**Fig. S1A**). DHT was not detected when islets were co-treated with the potent 5α-R inhibitors, finasteride and dutasteride (8) (**Fig. S1A**). Having observed that male mouse and human islets convert T to DHT, we next examined the physiological relevance of intracrine T metabolism to islet function in an experiment assessing GSIS in static incubation. At 16.7 mM glucose, male mouse islets exposed to T showed increased GSIS compared to those exposed to vehicle only (**Fig. S1B**). Notably, exposure of these islets to 5α-R inhibitors blocked the ability of testosterone to enhance GSIS (**Fig. S1B**). Together, these data demonstrate that male mouse islets convert T to DHT via 5α-R activity, a process essential to enhance GSIS.

Having observed that GLP-1 corrects the insulin secretory defect of cultured βARKO^RIP^ islets (**Fig. 1K**), we explored whether preventing GLP-1 degradation *in vivo* using linagliptin, a dipeptidyl peptidase-4 (DPP-4) inhibitor (9), would restore the β cell defect of male βARKO^MIP^ mice. After 4 weeks of linagliptin treatment, we studied βARKO^MIP^ mice and their littermate controls during IP (IP-GTT) and oral (O-GTT) glucose challenges. In response to O-GTT, linagliptin improved oral GSIS (**Fig. S1C**) and glucose tolerance (**Fig. S1D)** in both control and βARKO^MIP^ mice compared to those without linagliptin treatment. Notably, owing to the small number of mice in each group, linagliptin improvement of O-GSIS and O-GTT was significant only in the linagliptin-treated βARKO^MIP^ mice compared to vehicle-treated βARKO^MIP^ mice (**Fig. S1C**). While βARKO^MIP^ mice exhibited glucose intolerance following an IP-GTT, as previously shown, linagliptin-treated βARKO^MIP^ mice were no longer glucose intolerant compared to linagliptin-treated controls (**Fig. S1E**). Notably, at 2h into the IP-GTT, blood glucose was significantly lower in linagliptin-treated βARKO^MIP^ mice compared to untreated βARKO^MIP^ mice (**Fig. S1E**). Note that in cultured male mouse islets, linagliptin amplified GSIS compared to vehicle treatment, consistent with prolongation of islet derived-GLP-1 half-life. However, linagliptin did not amplify the insulinotropic effect of DHT on GSIS (**Fig. S1F**). Thus, prolonging GLP-1 half-life with linagliptin (which mostly affects circulating GLP-1) can improve the β cell response to a glucose load in absence of β cell AR.

Having established that DHT amplifies the insulinotropic effect of both islet-derived and exogenous GLP-1 via AR and GLP-1R in β cells in male mouse and human islets [**(Fig. 1)** and (2)], we sought to determine if DHT could similarly amplify the insulinotropic action of glucose-dependent insulinotropic polypeptide (GIP) and glucagon, which also bind class B G-protein coupled receptors (GPCR) coupled to Gα_S_ and adenylate cyclase (AC) (10). We examined the effect of DHT on cAMP production in 832/3 insulin-secreting cells, an incretin-responsive β cell model (11). We previously showed that DHT insulinotropic action is associated with increased islet cAMP production in an AR-dependent manner (2). To study DHT-induced cAMP production dynamically in live cells, we used an Epac-based FRET sensor (Epac2-camps) (12). In these conditions, GLP-1, GIP, glucagon, and the activator of AC, forskolin, all elicited a rapid and sustained rise in cAMP production compared to vehicle treated cells (**Fig. 2A-E**). Interestingly, DHT alone elicited a rise in cAMP production, although the onset of the cAMP increase was delayed compared to incretin treatments (**Fig. 2A-E**). Notably, when administered with GLP-1, DHT treatment elicited a further rise in GLP-1-induced cAMP production (**Fig. 2A-E**). The ability of DHT to amplify GLP-1 production of cAMP was also observed with the GLP1R agonist exendin 4 (**Fig. S2A**). In contrast, DHT had no effect on cAMP production induced by glucagon or GIP (**Fig. 2A-2E**). Further, DHT produced no effect on forskolin-induced cAMP production, arguing against DHT increasing the overall cellular capacity to produce cAMP via activation of AC. In fact, when forskolin was used at lower doses, DHT seemed to inhibit forskolin-induced cAMP production (**Fig. S2B**). Thus, DHT alone stimulates cAMP production and selectively enhances the ability of GLP-1 to produce cAMP (**Fig. 2A-E**). To assess the functional significance of DHT amplification of GLP-1-induced cAMP production, we next assessed the effect of DHT on GLP-1, GIP, glucagon and forskolin in an experiment assessing GSIS in static incubation in INS1 832/3 cells. The use of an insulin-secreting cell line in this experiment eliminates the confounding effect of islet-derived GLP-1, as well as paracrine effects mediated by glucagon itself. In these cells, DHT-alone did not significantly increase GSIS suggesting that the DHT-induced increase in cAMP described above (**Fig. 2A-E**) is insufficient to promote insulin vesicles exocytosis. However, consistent with cAMP production data (**Fig. 2A-E**), DHT amplified the insulinotropic effect of exogenous GLP-1 (**Fig. 2F**). In contrast, and also consistent with cAMP data, while both GIP and glucagon alone enhanced GSIS, DHT failed to amplify the insulinotropic effect of GIP (**Fig. 2G**) or glucagon (**Fig. 2H**). Further, forskolin also enhanced GSIS, but this effect was not amplified by DHT (**Figure 2I**). Importantly, similar results were obtained using wild-type male islets demonstrating that the selectivity of DHT for GLP-1 is observed in primary β cells (**Fig. S2C**). Thus, DHT selectively amplifies the insulinotropic action of GLP-1, but not that of GIP and glucagon.

**Figure 2.**
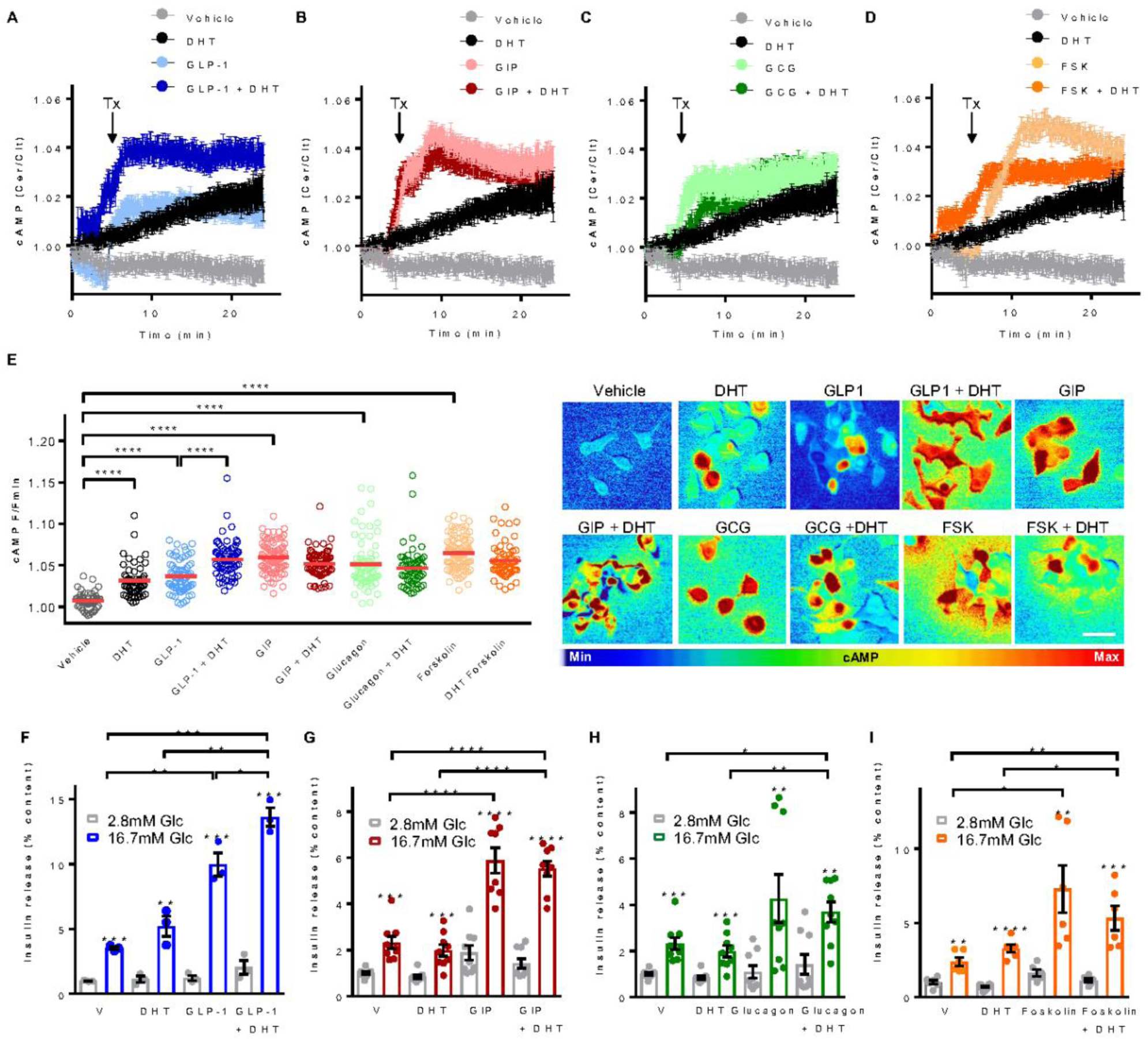
DHT amplifies the insulinotropic action of GLP-1 selectively. **(A-E)** 832/3 cells were infected with adenovirus harboring the FRET Epac2 camps probe and treated with DHT (10nM), **(A)** GLP-1 (10nM), **(B)** GIP (100nM), **(C)** glucagon (20nM) and **(D)** forskolin (FSK, 10uM) starting at the indicated time (Tx arrow, 5 min). cAMP production was monitored in real-time from live cells. Note that the same control trace is shown in (a-d), since all experiments were run in parallel but are shown on separate graphs for clarity. **(E)** Summary graph showing amplitude of cAMP responses from (A-C) of 6 independent experiments, each with 30-40 cells/treatment. Representative pictures are shown on the left. **(F-I)** GSIS in static incubation in 832/3 cells were treated with vehicle, DHT (10nM), **(F)** GLP-1 (10nM), **(G)** GIP (100nM), **(H)** glucagon (20nM) and **(I)** forskolin (100nM) for 40 minutes. Values represent the mean ± SE of n= 3 independent wells, and 2-4 independent experiments. * P < 0.05, ** P < 0.01, *** P < 0.001, **** P < 0.0001.

We wondered whether ligand-activated AR and GLP1R might interact within the same cellular compartment, as suggested by the selective effect of DHT on GLP1-, but not glucagon- and GIP-induced cAMP rises. To this end, we transfected INS1 832/3 cells with GLP-1R-GFP and FLAG-AR plasmids and studied receptor localization by immunofluorescence. At basal (vehicle), GLP-1R-GFP was localized to the plasma membrane and likely the Golgi compartments (due to receptor overexpression from a CMV promoter in the plasmid). FLAG-AR was localized close to the plasma membrane and throughout the cytosol (**Fig. 3A and Fig. S3**). Note that, as a result of AR overexpression, a pool of FLAG-AR was also localized to the nucleus, which is not observed when staining the endogenous AR (2). DHT treatment did not alter the location of FLAG-AR or GLP-1R-GFP. Following GLP-1 stimulation alone or in combination with DHT, a fraction of internalized GLP-1R-GFP came in proximity to the FLAG-AR fraction, but obvious co-localization was not detected (**Fig. 3A and Fig. S3**). Indeed, following staining of early and recycling endosomes with transferrin, we determined that stimulation with GLP-1 alone or with DHT triggered GLP-1R-GFP internalization to an endocytic location (**Fig. S4**), but AR did not localize to endosomes under any conditions (**Fig. S5**). Together these data suggest that, upon GLP-1 stimulation, a fraction of GLP-1R internalizes to endosomes where it comes in proximity with a pool of DHT-activated AR. However, AR and GLP-1R do not directly interact in the same cellular micro domain.

**Figure 3.**
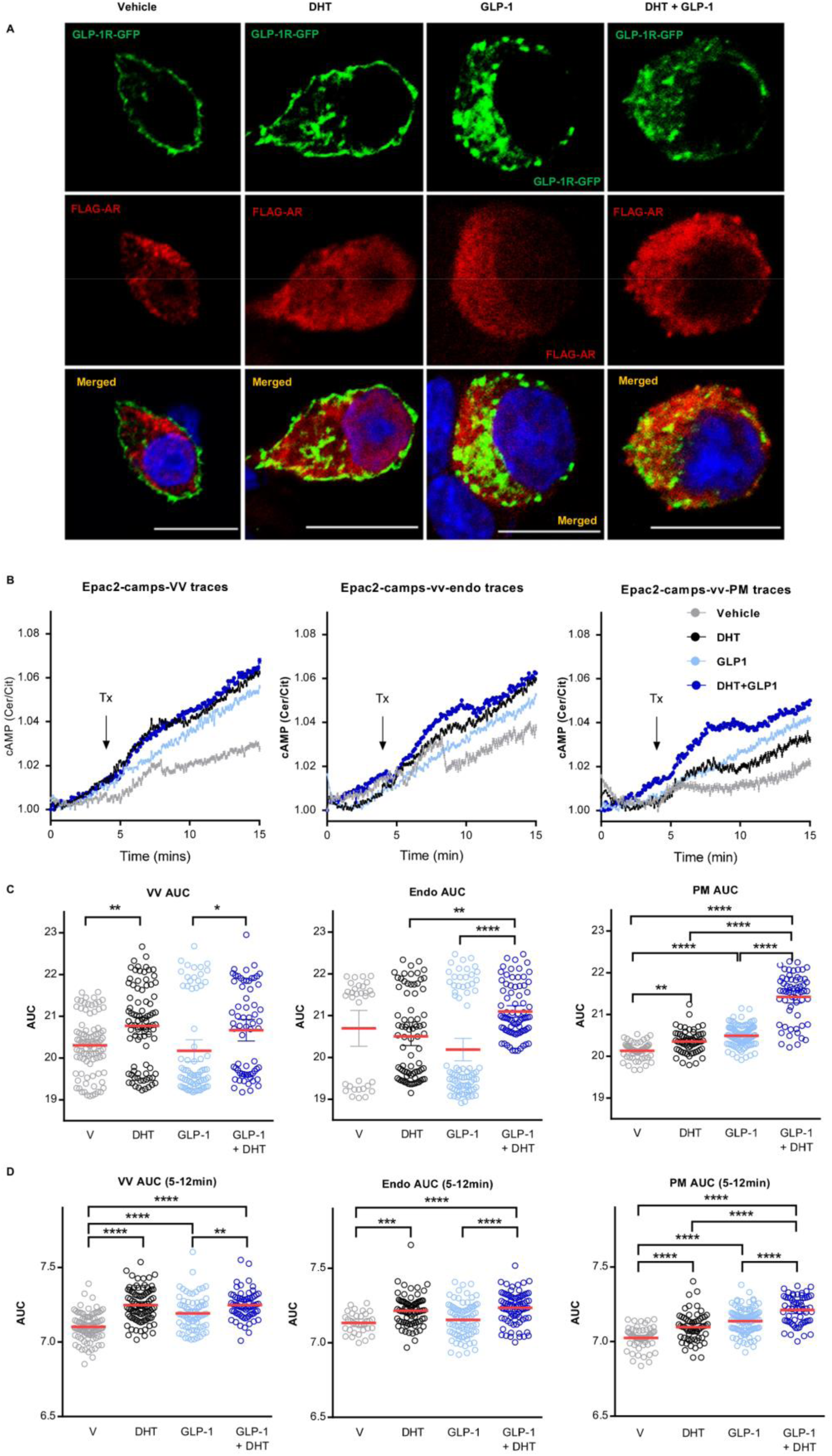
DHT amplifies cAMP production at the plasma membrane and endosomes. **(A)** INS1 832/3 cells were transfected with GLP-1R-GFP (green) and FLAG-AR (red) and treated with vehicle, DHT (10nM), GLP-1 (10nM) and DHT + GLP-1 for 15 min. Receptor localization was assessed by immunofluorescence. **(B-D)** 832/3 cells were transfected with organelle-targeted cAMP FRET biosensors and treated with DHT (10nM), GLP-1 (10nM) and DHT + GLP-1. Epac2-camps plasmids targeted to the cytoplasm (Epac2-camps-VV), the endosomes (Epac2-camps-vv-Endo) or the plasma membrane (Epac2-camps-vv-PM) were used. **(B)** cAMP production was monitored in real-time from live cells with treatment starting at the indicated time (Tx arrow, 5 min). **(C)** Total cAMP area under the curve (AUC), **(D)** AUC of the cAMP peak between 5- and 12-min. Values represent the mean ± SE. * P < 0.05, ** P < 0.01, *** P < 0.001, **** P < 0.0001.

Upon ligand binding, endosomal trafficking of the GLP-1R promotes compartmentalized endosomal cAMP generation from endosomally-associated AC, which is important for promoting insulin granule exocytosis (13, 14). However, the biased agonist exendin-phe1, which promotes membrane retention of the GLP-1R, also increases cAMP production and GSIS, particularly in the long term following continued agonist exposure (15). Therefore, both GLP-1R membrane and endosomal cAMP production seem important for GSIS. To determine if DHT enhances GLP-1-induced cAMP production by altering the trafficking of the GLP-1R, we studied the effect of DHT in MIN6B1 cells (a subclone of mouse insulinoma MIN6 cells with enhance insulin secretory responsiveness) stably expressing SNAP-tagged GLP-1R following treatment with either exendin-4 (which promotes GLP-1R internalization) or exendin-phe1 (which promotes GLP-1R membrane retention) (15). We did not detect any effect of DHT on GLP-1R trafficking when cells were pre-treated with exendin-4 (**Fig. S6A**) or exendin-phe1 (**Fig. S6A**), suggesting that DHT does not perturb GLP-1R membrane trafficking but rather enhances GLP-1R signaling from membrane or intracellular locations.

Upon binding of GLP-1 to its receptor, the GLP1R-ligand complex triggers cAMP signal at the plasma membrane and at endosomes, which is critical to promote insulin vesicle exocytosis.(13-15) To determine if DHT enhances cAMP production in these compartments, we generated new cAMP sensors specific to the plasma membrane (_T_Epac_VV_-PM) and endosomes (_T_Epac2_VV_-Endo) from the cytoplasmic _T_Epac_VV_ biosensor, also used by us as a control (**Fig. S6B**), or cytoplasm (control, Epac2-camps-VV). Using these organelle-specific cAMP sensors, we studied GLP-1 and DHT-induced subcellular production of cAMP in INS1 832-3 cells. Compared to vehicle, both GLP-1 and DHT increased cAMP production in cytoplasm, at the plasma membrane and the endosomes (**Fig. 3B and C**). Importantly, DHT amplified GLP-1-induced cAMP production predominantly at the plasma membrane and endosomal compartments with a maximum effect in the first 5 minutes (**Fig. 3B and D**), consistent with the trafficking analysis.

We reasoned that, if DHT amplifies GLP-1-induced cAMP production at the plasma membrane and endosomal compartments, DHT should sensitize cAMP-dependent pathways that promote insulin vesicles exocytosis in these compartments, namely, the protein kinase A (PKA) and the exchange protein activated by cAMP islet 2 (Epac2) pathways (16). Indeed, cAMP elevation causes rapid binding of Epac2A to the β cell plasma membrane, where it accumulates at secretory granules and sensitizes them to undergo exocytosis (17, 18). We examined DHT-induced amplification of GSIS in static incubation in wild-type male mouse islets following pharmacological inhibition of PKA and Epac2. In these conditions, DHT potentiated GSIS in male mouse islets and both the PKA inhibitor H89 (**Fig. 4A**) and the Epac2 inhibitor ESI-09 abolished this effect (**Fig. 4B**). Further, in islets from male human donors, the ability of DHT to amplify GSIS was similarly abolished following inhibition of PKA and Epac2 (**Fig. 4C and D**).

**Figure 4.**
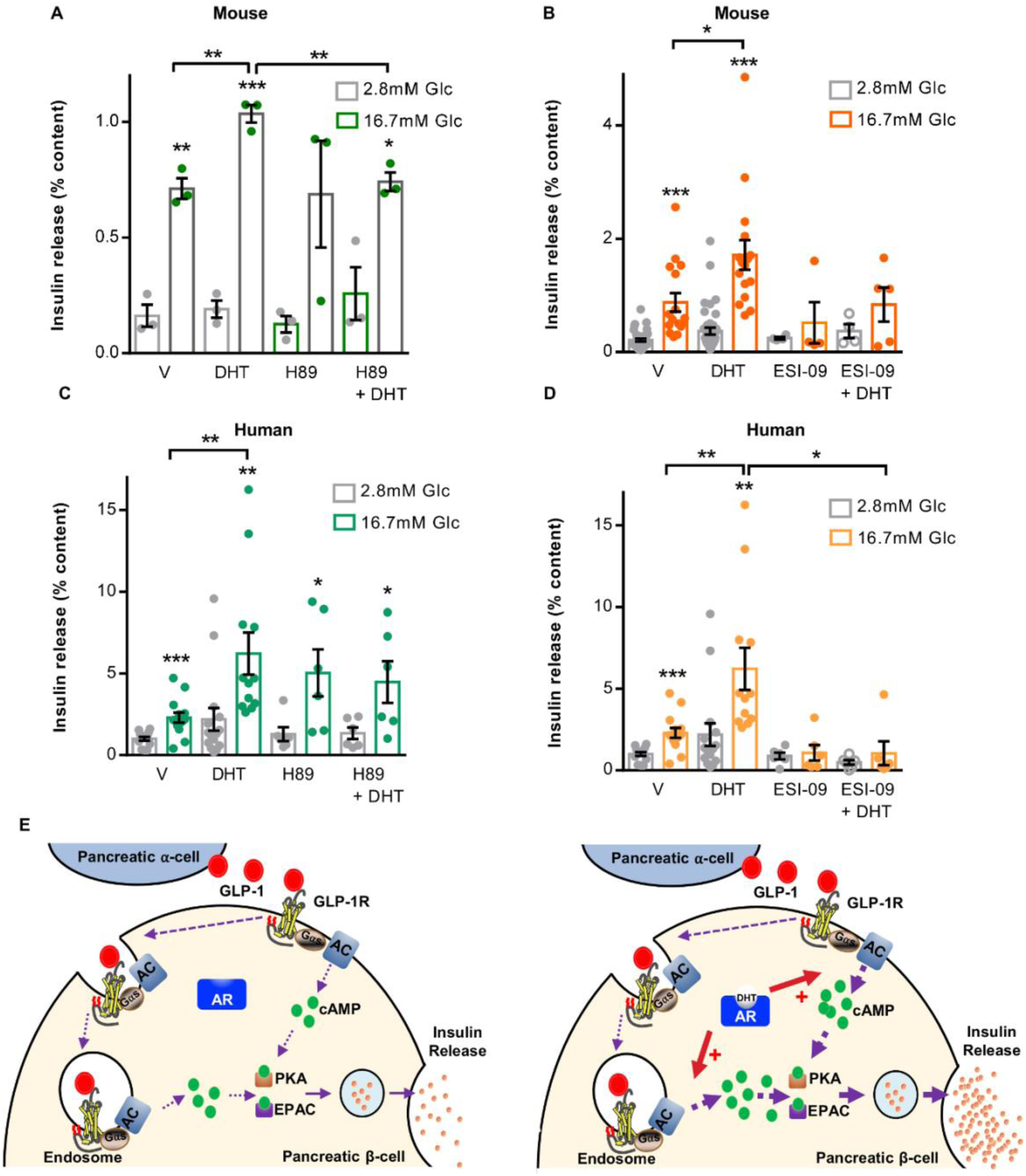
DHT insulinotropic action requires PKA and EPAC in mouse and human islets. GSIS was measured in static incubation in **(A, B)** wild-type male mouse and **(C, D)** male human islet donors treated with vehicle, DHT (10nM) for 40 minutes in the presence of absence of **(A, C)** H89 (10uM) or **(B, D)** ESI-09 (10uM). Values represent the mean ± SE of n= 2 mice/group and 2 human islet donors measured in triplicate. * P < 0.05, ** P < 0.01, *** P < 0.001. **(E)** Proposed mechanism of DHT-enhanced GLP-1-induced cAMP production at the plasma membrane and endosomes in β-cells, leading to PKA and EPAC activation and increased insulin secretion.

## Discussion

This study reveals that testosterone activation of AR in β cells amplifies GLP-1R-mediated production of cAMP at the plasma membrane and endosomal compartments, activating PKA and Epac2 to promote insulin vesicles exocytosis, and these pathways are conserved in human β cells as summarized in **Fig. 4E**. Compartmentalization of cAMP signaling microdomains within β cells permits the second messenger to mediate various independent cellular processes. Ligand-activated GLP-1R compartmentalizes into endosomal membranes in endosomes to produce cAMP from endosomally-associated AC (14). DHT activation of AR does not seem to alter the trafficking of the GLP-1R. Instead, using novel cAMP probes, we show that DHT alone enhances cAMP production in these compartments. However, DHT alone is insufficient to sensitize pools of insulin granules for exocytosis and promote GSIS. Notably, DHT amplifies the ability of GLP-1 to produce cAMP at the plasma membrane and endosomes. Accumulating evidence supports a model in which internalization of GPCRs in endosomal compartments allows cAMP initiated at the plasma membrane to be prolonged in the endosomal station (19-21). At the same time, agonist-mediated prolonged plasma membrane retention of the GLP-1R increases cAMP production and GSIS (15). Here, the cumulative effects of DHT and GLP-1 on cAMP generation at both the plasma membrane and endosomal compartments produces a massive sensitization of the releasable pool of insulin granules for exocytosis (13, 14). Consistent with this finding, DHT enhancement of GSIS in male mouse and human islets relies on PKA and Epac2 activity, which are located at the plasma membrane and endosomes, and instrumental in insulin granule exocytosis.

An important unanswered question relates to the mechanism by which ligand-activated AR increases cAMP production to selectively sensitize GLP-1R. There is little evidence of co-localization of AR and GLP-1R under any condition we tested. The effect of AR on GLP-1R production of cAMP is likely to be indirect, perhaps by modulating the activities of GαS or ACs. GLP-1 and glucagon receptors have been reported to elicit a rise in cAMP and promote GSIS via differential stimulation of membrane-anchored AC (mAC) (22), and soluble AC (23). DHT is unlikely to increase the activity of mAC as it shows no effect on cAMP production and GSIS induced by the activator of mAC, FSK. Alternatively, GLP-1, GIP, and glucagon-regulated cAMP micro domains could be spatially distinct. Further studies are needed to address these important issues.

In conclusion, ligand-bound AR and GLP1-R exert synergistic effects on cAMP at the plasma membrane and endosomes to enhance GSIS. This study establishes a novel biological paradigm in which membrane location of a steroid nuclear receptor enhances the ability of a G protein-coupled receptor to produce cAMP.

## Acknowledgments

This work was supported by grants from the National Institutes of Health (DK107444 and DK074970), a U.S. Department of Veterans Affairs Merit Award (BX003725) and an Investigator-Initiated Study from Boehringer Ingelheim and Eli Lilly to FMJ. WA was supported by a Wellcome Trust Investigator Award (209492/Z/17/Z), and the National Institute for Health Research (NIHR) Birmingham Biomedical Research Centre at the University Hospitals Birmingham NHS Foundation Trust and the University of Birmingham (Grant BRC-1215-20009). D.J.H. was supported by MRC (MR/N00275X/1 and MR/S025618/1) and Diabetes UK (17/0005681) Project Grants. This project has received funding from the European Research Council (ERC) under the European Union’s Horizon 2020 research and innovation programme (Starting Grant 715884 to D.J.H.). Human islets were provided by the Integrated Islet Distribution Program (IIDP) funded by the National Institute of Diabetes and Digestive and Kidney Diseases and with support from the Juvenile Diabetes Research Foundation International.

## Author Contributions

WX designed and performed experiments including metabolic studies in mice, GSIS in cultured mouse and human islets, analyzed the data, prepared the figures, wrote and edited the manuscript. FBA performed experiments of dynamic measurement of cAMP in live cells. SB performed experiments of GLP-1R and AR co-localization in INS1 832/3 cells and SNAP-GLP-1R internalization in MIN6B1 cells. LS performed steroid conversion experiments including androgen profiling by UHPLC-MS/MS. MMFQ extracted images of AR and GLP-1R localization and prepared all figures of IHC. WA designed and analyzed the data of UHPLC-MS/MS. AT designed and analyzed experiments of GLP-1R and AR co-localization in INS1 832/3 cells and SNAP-GLP-1R internalization in MIN6B1 cells. DJH designed and analyzed experiments of dynamic measurement of cAMP in live cells. FMJ designed the study, analyzed the data, wrote and revised the manuscript. All authors reviewed and edited the manuscript and accepted the final version.

## Materials and Methods

### Mutant mice

To generate βARKO^MIP^ mice we crossed mice carrying the AR gene with floxed exon 2 on their X chromosome (AR^lox^) with the Ins1-Cre/ERT (MIP-Cre^+/-^) transgenic mouse (Jackson Lab). Generation and characterization of ARlox ^-/-^ have been described (24). We induced tamoxifen (tam; Sigma) inactivation of AR after puberty and following 4 weeks of tam treatment in silastic tubing, and all of metabolic measures were taken after 4-week waiting period. 10mm silastic laboratory tubing (Dow Corning) was filled with 15mg tam, capped with wooden applicator sticks, and sealed with silastic medical adhesive (Dow Corning). The βARKO^RIP^ mouse was generated by crossing AR^lox^ with transgenic mice overexpressing the Cre recombinase under control of the RIP promoter (RIP-Cre, Jackson Laboratory). The βGLP-1KO^RIP^ mouse was generated by crossing GLP-1R^lox^ (a kind gift from Dr. David A. D’Alessio of Duke University) (25) with RIP-Cre. All mice were generated on a C57/BL6 background. All studies were performed with the approval of the Tulane University Animal Care and Use Committee in accordance with the NIH Guidelines.

### Western Diet

Mice were weaned onto a customized diet designed to be high in saturated fat and simple sugars (sucrose and fructose) to mimic a western diet (30% AMF; 14.9% Kcal protein, 33.2% Kcal carbohydrates, 51.9% Kcal fat; Harlan Teklad) for 9 weeks.

### Metabolic studies

Blood glucose was measured from tail vein blood using True Metrix (Trividia Health). Insulin was measured in plasma by ELISA kit (Millipore). For IP-GTT (2 g/kg) and GSIS (3 g/kg), mice were fasted overnight before glucose injection.

### Mouse islet isolation and insulin secretion in static incubation

Islet isolation was performed following pancreatic duct injection with collagenase (Sigma) as described.(26) For measurement of insulin secretion, islets were hand-picked under a dissection microscope and treated with DHT (10nM; Steraloids), testosterone (10nM; Sigma), finasteride (100nM; Sigma), dutasteride (100nM; Sigma), GLP-1 (10nM; kind gift from Dr. Richard DiMarchi), GIP (100nM; Tocris), glucagon (20nM; Sigma), forskolin (10uM; Sigma), H89 (10uM; CST), ESI-09 (10uM; Sigma), or vehicle (95% ethanol) at 2.8mM and then 16.7mM glucose for 40 minutes sequentially. Insulin release from islets was measured as described with Rat/Mouse Insulin ELISA kit (Millipore Sigma) (26).

### Measurement of insulin secretion in static incubation in human islets

Human islets were hand-picked under a dissection microscope, and treated with DHT (10nM; Sigma), H89 (10uM; CST), ESI-09 (10uM; Sigma), or vehicle (95% ethanol) at 2.8mM and then 16.7mM glucose for 40 minutes sequentially. Insulin release from islets was measured with Human Insulin ELISA kit (Millipore Sigma). See **Table S1** for donor information.

### Generation of organelle-targeted FRET cAMP biosensors

Organelle-targeted cAMP FRET biosensors were generated from the untargeted Epac- and mTurquoise-based _T_Epac_VV_ sensor (a gift from Prof Kees Jalink, The Netherlands Cancer Institute)(27) as follows: Endo-_T_Epac_VV_ was generated by PCR cloning of the 2xFYVE domain (endosomal targeting sequence) of the bPAC-Endo construct (a kind gift from Prof Mark Von Zastrow, UC San Francisco)(21) to the N-terminus end of _T_Epac_VV_ using HindIII and NotI restriction sites. PM-_T_Epac_VV_ was generated by PCR cloning of the lipid raft-binding domain of Lyn kinase (plasma membrane targeting sequence) of the bPAC-PM construct to the N-terminus end of _T_Epac_VV_ using the same restriction sites as above. bPAC-Endo and bPAC-PM construct were a gift from Prof Mark Von Zastrow, UC San Francisco (21).

### Immunofluorescence analysis of GLP-1R and AR co-localization

INS1 832/3 cells were plated in 24-well plates and the following day, transfected with GLP-1R-GFP and FLAG-AR plasmids (0.5μg of each plasmid/well plus 2μL Lipofectamine 2000). After 24h, the cells were trypsinized and plated in 24-well plates with glass coverslips using phenol-red-free medium containing charcoal-stripped FBS (RPMI 1640 no phenol red, 10% v/v charcoal-stripped FBS, 100 units/mL penicillin, 100 μg/mL streptomycin, 10mM HEPES, and 1mM sodium pyruvate). 48h post-transfection, the cells were treated with transferrin-555 for 30 minutes to label the endosomal pathway. Subsequently, treatments with control medium, GLP-1 100nM, DHT 10nM, or GLP-1 + DHT for 5 or 10 minutes were performed. After a wash with ice-cold PBS, the cells were fixed in 4% PFA for 20 minutes at 4C. After two washes with PBS, 0.1% Triton X-100 was added for 10 min to allow permeabilization. Cells were washed with PBS twice and blocking buffer (3% w/v BSA, 1% v/v goat serum, 0.1% v/v Tween-20 in PBS) was added for 30 minutes. The blocking buffer was discarded and primary anti-FLAG antibody (mouse monoclonal antibody, F3165 Sigma) diluted 1:500 in blocking buffer was added overnight at 4C. After three 5-minute washes in PBS + 0.1% v/v Tween-20 (PBST), secondary antibody anti-mouse Alexa Fluor 647 (1:1,000 in blocking buffer) was incubated for 30 minutes at room temperature. Following three 5-minute washes in PBST, the coverslips were mounted onto slides using ProLong Diamond Antifade Mountant with DAPI (ThermoFisher Scientific). Images were captured using a Zeiss LSM-780 inverted confocal microscope with a 63x/1.4numerical aperture oil-immersion objective from the Facility for Imaging by Light Microscopy (FILM) at Imperial College London, and analyzed in Fiji.

### Dynamic measurement of cAMP production

INS-1 832/3 cells were cultured in 10ml complete medium composed of RPMI 1640 medium supplemented with 10% fetal calf serum, 100 IU/ml penicillin, 100 mg/l streptomycin, 10 mM HEPES, 2 mM L-glutamine, 1 mM sodium pyruvate, and 50 µM beta-mercaptolethanol. Cells were split twice a week and cultured in a 37°C incubator containing 5% CO2. For imaging INS-1 832/3 cells were grown on glass coverslips. Adenoviral infection with Epac2-camps was performed 48h prior to imaging. Transfection of INS-1 832/3 cells with Epac2-camps plasmids was performed using Lipofectamine-2000 with 1 µg DNA 24h prior to imaging. Imaging of cAMP was conducted using a Crest X-Light spinning disk, coupled to a Nikon Ti-E, SPECTRA X light engine and 20 x objective, as described (12). cAMP responses were measured using adenovirus harboring the FRET probe Epac2-camps (a kind gift from Prof Dermot Cooper, University of Cambridge), or alternatively Epac2-camps plasmids targeted to the plasma membrane (PM-_T_Epac_VV_), cytoplasm (_T_Epac_VV_) or endosomes (Endo-_T_Epac_VV_) (see above for details on their cloning). Excitation was delivered at λ = 430-450 nm, with emitted signals detected at λ = 460-500 nm and 520-550 nm for Cerulean and Citrine, respectively. HEPES-bicarbonate buffer was used, containing (in mM): 120 NaCl, 4.8 KCl, 24 NaHCO_3_, 0.5 Na_2_HPO_4_, 5 HEPES, 2.5 CaCl_2_, 1.2 MgCl_2_, 16.7 *D*-glucose. FRET responses were calculated as the fluorescence ratio of Cerulean/Citrine and normalized as F/Fmin, where R denotes fluorescence at any given time point and R_0_ denotes fluorescence at time 0.

### Confocal analysis of SNAP-GLP-1R internalization

MIN6B1 cells (a gift from Prof Philippe Halban, University of Geneva) stably expressing human SNAP-GLP-1R (“MIN6B1-SNAP-GLP-1R”)(15) were generated by transfection of a SNAP-GLP-1R vector (Cisbio) followed by G418 selection and maintained in DMEM with 15% FBS, 50 µM β-mercaptoethanol and 1% penicillin/streptomycin. Cells were seeded in 24-well plates with glass coverslips using medium containing charcoal-stripped FBS (10% v/v). 24h later, cells were labelled at 37C for 30 min with 1 µM SNAP-Surface 549 to label surface receptors, treated for 30 min with 100 nM Exendin-4 or the biased agonist Exendin-Phe1(15) in the presence or absence of 10 nM DHT. After a wash with ice-cold PBS, the cells were fixed in 4% PFA for 20 minutes at 4C, washed in PBS, mounted onto slides using ProLong Diamond Antifade Mounting Medium with DAPI (ThermoFisher Scientific). Images were captured using a Zeiss LSM-780 inverted confocal microscope with a 63x/1.4numerical aperture oil-immersion objective from the Facility for Imaging by Light Microscopy (FILM) at Imperial College London, and analyzed in Fiji.

### Statistical analysis

Statistical analyses were performed with GraphPad Prism. Results are presented as mean ± SEM as specified in figures. When results showed a Gaussian distribution, statistical analyses were performed using the unpaired Student’s *t* test or ANOVA (with Bonferroni post hoc test). A P value less than 0.05 was considered statistically significant. * P<0.05, ** P<0.01, ***P < 0.001, ****P < 0.0001.

