## Supplemental Data for "Testosterone enhances GLP-1 efficacy at the plasma membrane and endosomes to augment insulin secretion in male pancreatic β cells"

### Supplemental Information

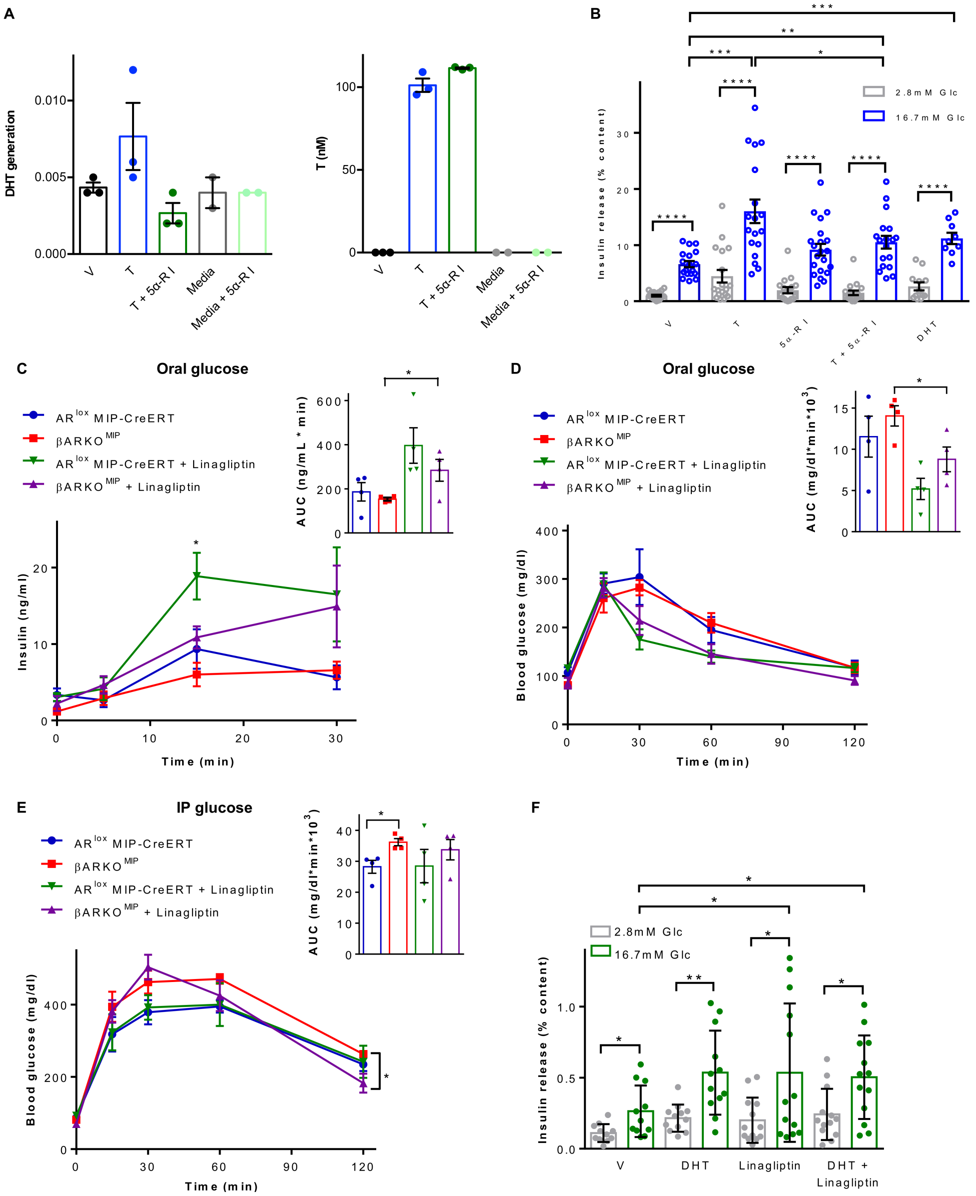


**Figure S1. (A)** Steroid quantification by ultra-high-performance liquid-chromatography tandem mass spectrometry (UHPLC-MS/MS). Generation of DHT from T (100nM) in male mouse islets is blocked by 5α-reductase inhibitors (5α-RI), finasteride (100nM) and Dutasteride (100nM). DHT concentrations were <LLOQ (0.24 nM) and hence no accurate quantification could be performed. DHT generation is shown as the ratio of the peak area of DHT and the peak area of its internal standard DHT-d3. Right: T recovery in the media. Values represent the mean ± SEM and scatter plot of technical triplicates for n=1 experiment (islets pooled from 10 male mice). **(B)** GSIS was assessed in static incubation in cultured male islets from C57/BL6 mice. Cultured islets were treated with vehicle, DHT (10nM), the finasteride (10nM) and dutasteride (10nM), for 40 minutes prior to measurement of insulin release by ELISA. **(C-E)** Mice were exposed to a western diet since weaning and linagliptin was added to the diet (83mg/kg of diet) 4 weeks before investigations. **(C)** Oral-GSIS (3 g/kg) with insulin area under the curve (AUC), **(D)** Oral-GTT (2 g/kg) with glucose AUC and **(E)** IP-GTT (2 g/kg) with glucose AUC were performed in the indicated mice with and without linagliptin treatment. *P < 0.05, βARKO^MIP vs. βARKO^MIP+ linagliptin at 120 min. Mice were studied at 40-45 weeks of age **(F)** GSIS was performed in static incubation in wild type C57BL/6 male mouse islets treated with vehicle, DHT (10nM), linagliptin (50nM) or DHT plus linagliptin for 40 minutes. Values represent the mean ± SE of n= 3-5 mice/group measured in triplicate. *P < 0.05, **P < 0.01, ***P < 0.001, ****P < 0.0001.

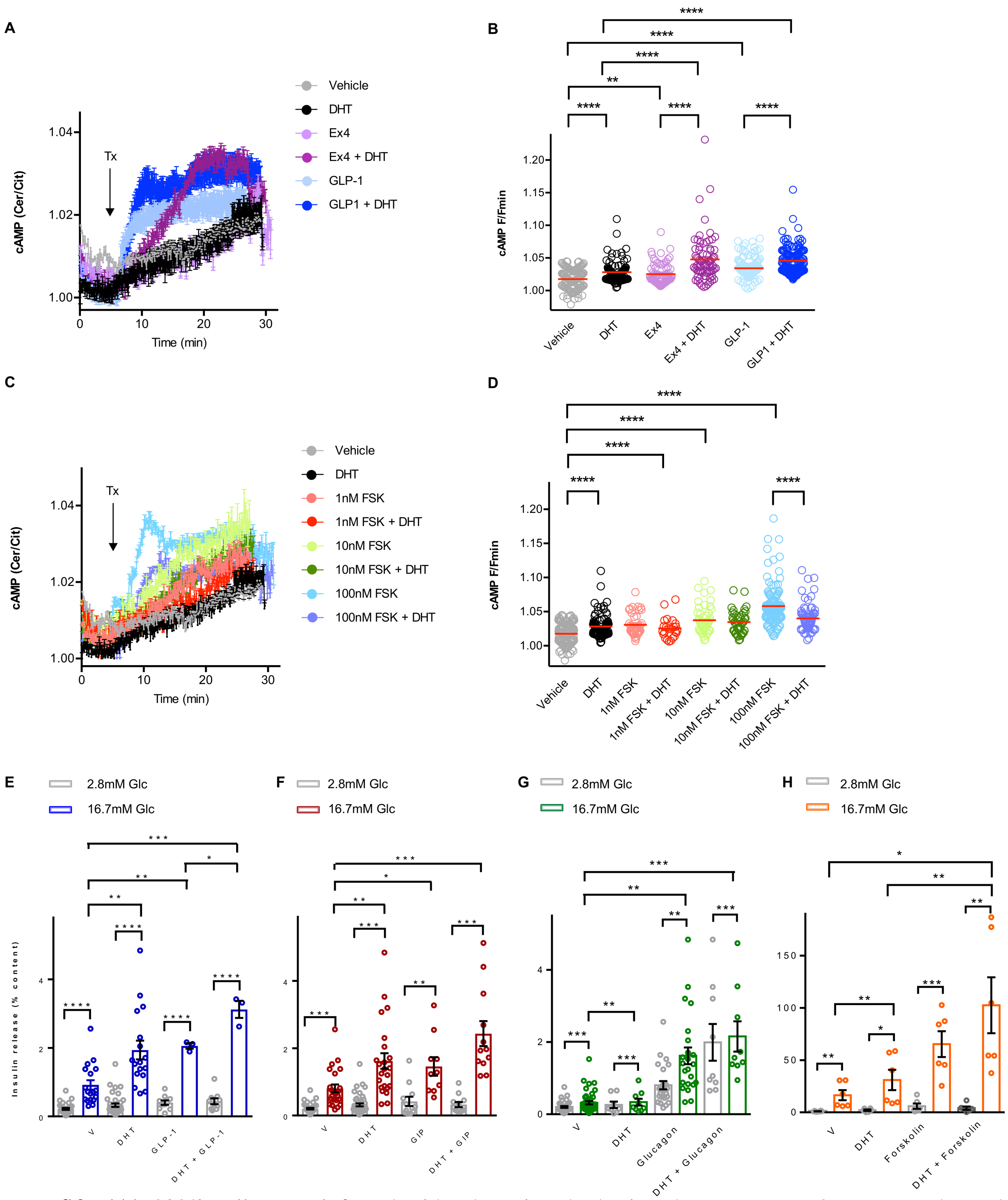


**Figure S2.** **(A)** 832/3 cells were infected with adenovirus harboring the FRET Epac2 camps probe and treated with DHT (10nM), GLP-1 (10nM), and exendin4 (Ex4, 10nM) starting at the indicated time (Tx arrow, 5 min). cAMP production was monitored in real-time from live cells. **(B)** Summary graph showing amplitude of cAMP responses from **(A)**. **(C)** 832/3 cells were infected with adenovirus harboring the FRET Epac2 camps probe and treated with DHT (10nM) and forskolin (FSK) at the indicated concentration starting at the indicated time (Tx arrow, 5 min). cAMP production was monitored in real-time from live cells. **(D)** Summary graph showing amplitude of cAMP responses from **(C)**. Note that vehicle/DHT values in **(A)** and **(C)** are identical, since the Ex4, GLP1 and FSK states were run in parallel with the same controls, but are shown on separate graphs for clarity. GSIS was assessed in static incubation in cultured male islets from C57/BL6 mice. Cultured islets were treated with vehicle, DHT (10nM), and **(E)** GLP-1 (10nM), **(F)** GIP (100nM), **(G)** glucagon (20nM) and **(H)** forskolin (100nM) for 40 minutes prior to measurement of insulin release by ELISA. Values represent the mean ± SE of n= 2-4 mice/group measured in triplicate. Values represent the mean ± SE. *P < 0.05, **P < 0.01, ***P < 0.001, ****P < 0.0001

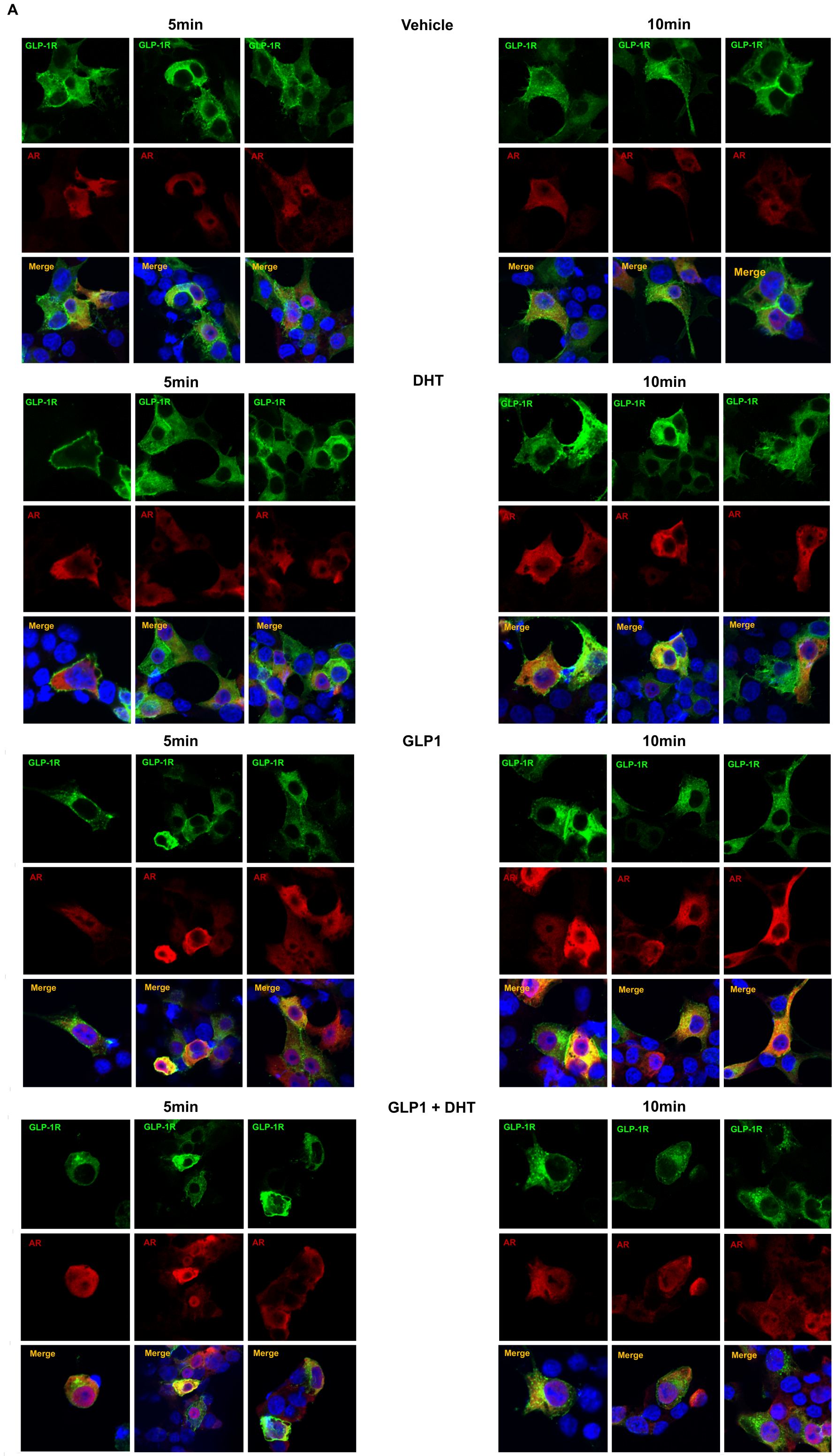


**Figure S3. (A)** INS1 832/3 cells were transfected with GLP-1R-GFP (green) and FLAG-AR (red) and treated with vehicle, DHT (10nM), GLP-1 (10nM) and DHT + GLP-1 for 5 or 10 min. Receptors localization was assessed by immunofluorescence. Images were captured by confocal microscopy. Representative pictures are shown.

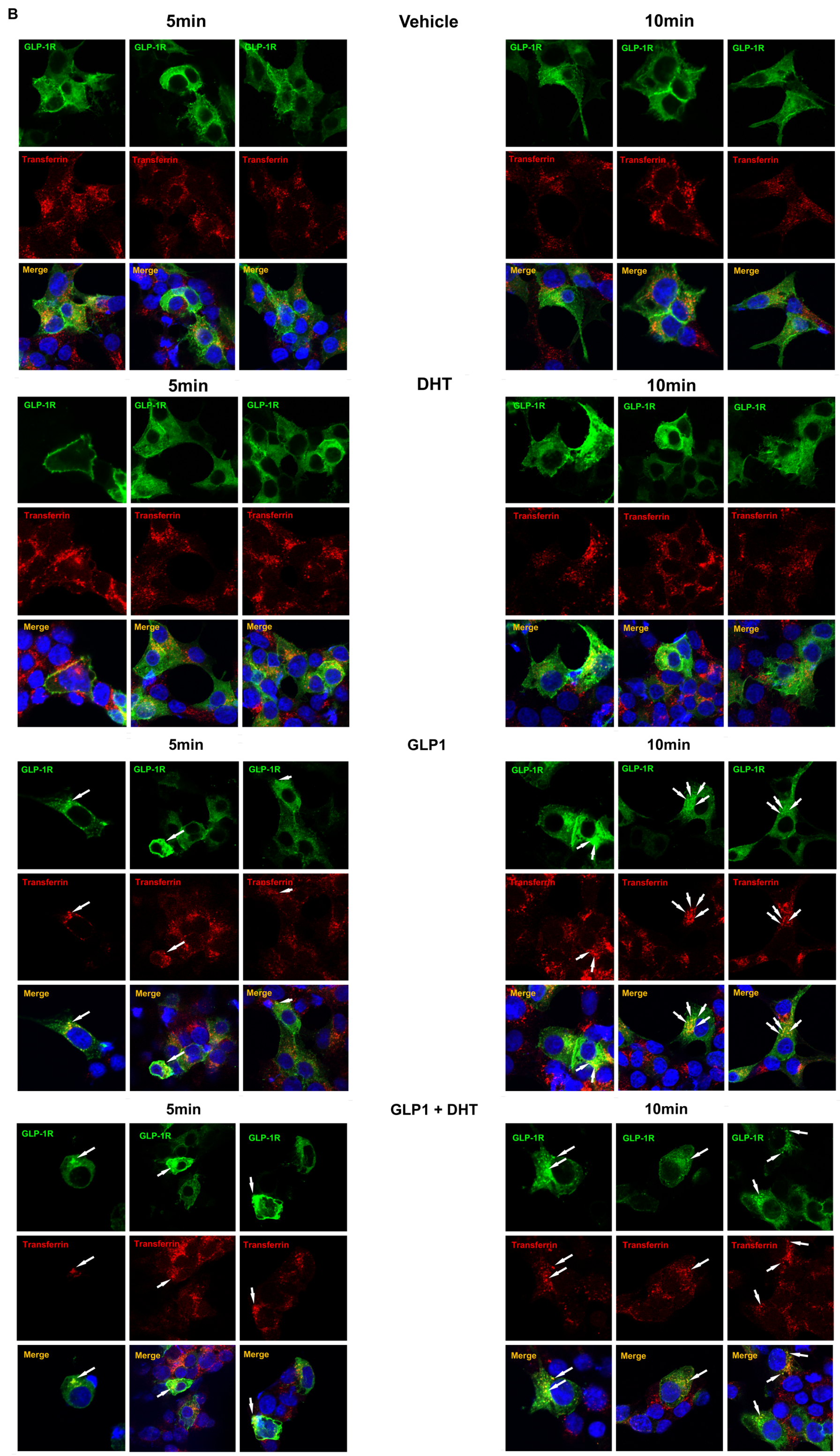


**Figure S4. (B)** INS1 832/3 cells were transfected with GLP-1R-GFP (green) and FLAG-AR (red) and treated with vehicle, DHT (10nM), GLP-1 (10nM) and DHT + GLP-1 for 5 or 10 min. Cells were treated with transferrin-555 to label the endosomal pathway. GLP-1R colocalization with endosomes was assessed by immunofluorescence and is shown with white arrows. Images were captured by confocal microscopy. Representative pictures are shown.

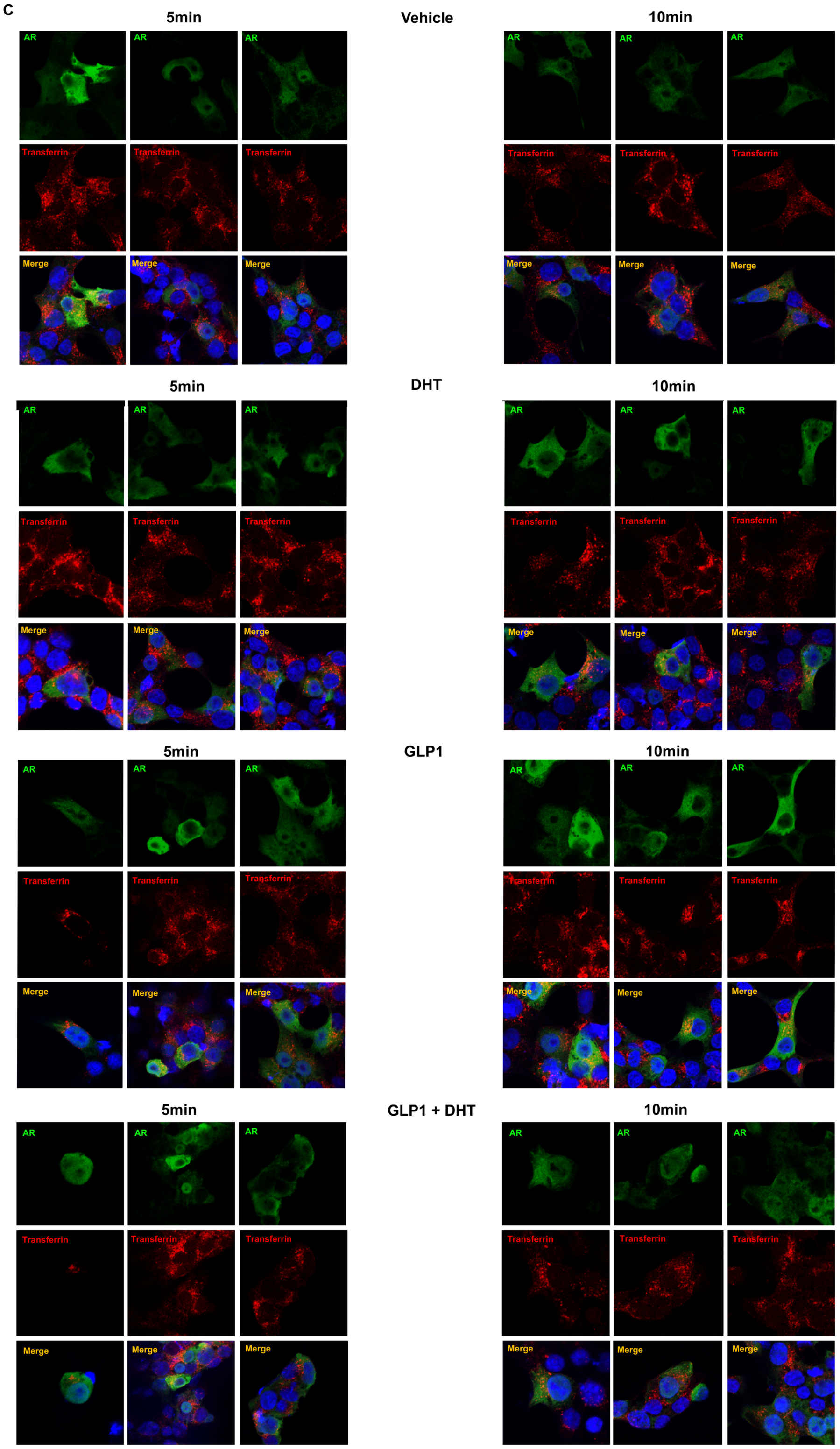


**Figure S5. (C)** INS1 832/3 cells were transfected with GLP-1R-GFP (green) and FLAG-AR (red) and treated with vehicle, DHT (10nM), GLP-1 (10nM) and DHT + GLP-1 for 5 or 10 min. Cells were treated with transferrin-555 to label the endosomal pathway. AR colocalization with endosomes was assessed by immunofluorescence and AR does not localize in endosomes. Images were captured by confocal microscopy. Representative pictures are shown.

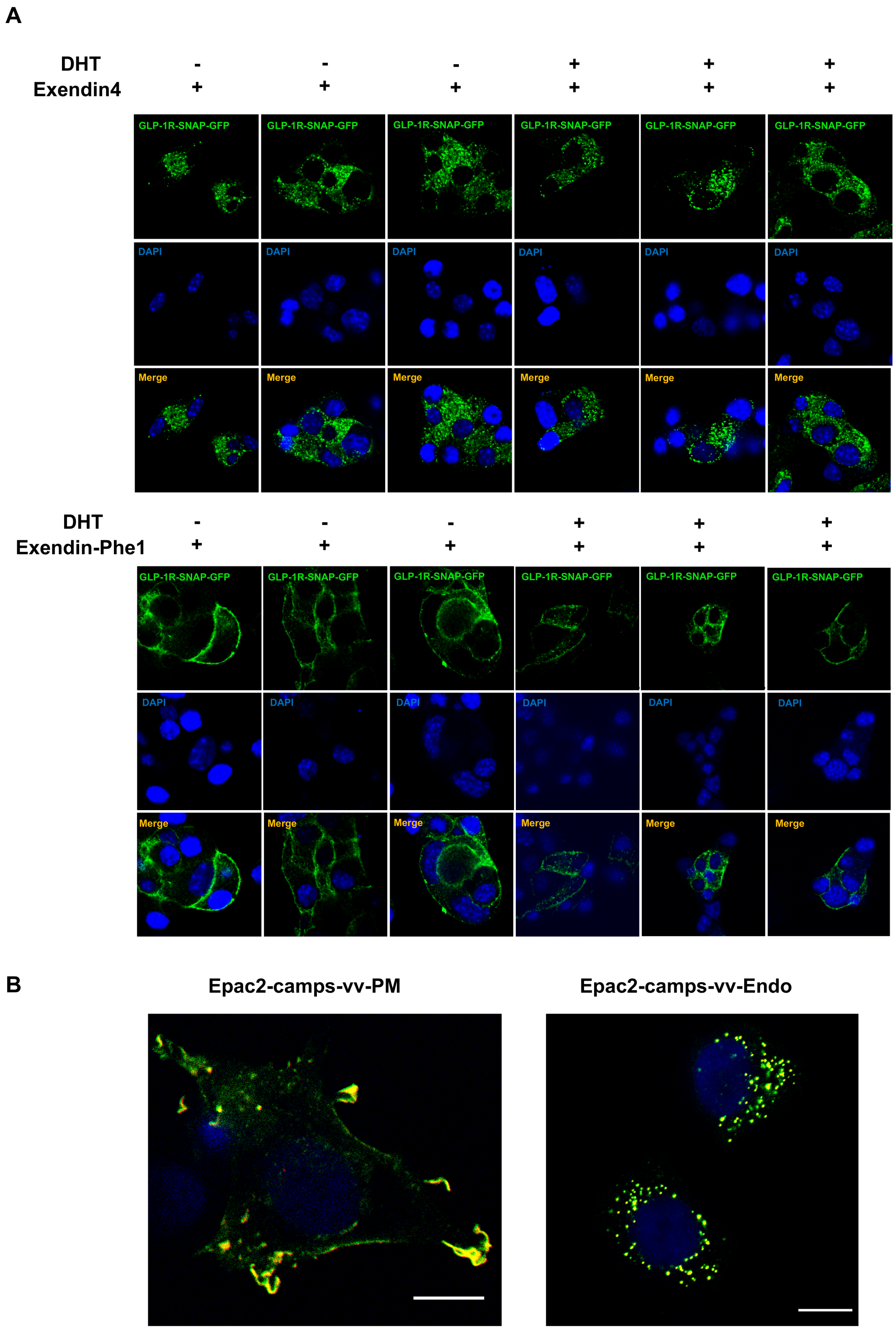


**Figure S6. (A)** MIN6B1 cells stably expressing human SNAP-GLP-1R cells were labelled with SNAP-Surface 549 to label surface receptors and treated for 30 min with 100 nM Exendin-4 or the biased agonist Exendin-Phe1 in the presence or absence of 10 nM DHT. Images were captured by confocal microscopy. Representative pictures are shown. **(B)** Chinese hamster ovarian (CHO-K1) cells were transfected with cAMP biosensors, Epac2-camps-vv-PM (plasma membrane localized) or Epac2-camps-vv-Endo (endosomally localized) and images were captured by confocal microscopy 24 hours after transfection. Note peripheral distribution for the plasma membrane biosensor and punctate pattern for the endosomal one. The scale bar is 10μM.

**Table S1. Human Islet Donor Profile**

| # | Receipt Date | Sex | Age (years) | Race | BMI | Cause of death | Center |
|---|---|---|---|---|---|---|---|
| 1 | Nov-18 | M | 44 | White | 40 | Stroke | South California Islet Cell Resources Center |
| 2 | Nov-18 | M | 58 | N/A | 32.4 | Head trauma | University of Wisconsin |

Table S2. Mass-to-charge (m/z) Transitions Used for the Quantification of Androgens

| Steroid | *m/z* transitions Quantifier Qualifier |
|---|---|
| **Analytes** | |
| Testosterone | 289.1 > 96.9 |

| | |
|---|---|
| | 289.1 > 109.0 |
| Dihydrotestosterone (DHT) | 291.3 > 255.1<br>291.1 > 159.0 |
| **Internal standards** | |
| Testosterone-D3 | 292.1 > 96.9 |
| DHT-D3 | 294.1 > 258.1 |

### Supplemental Experimental Procedures

**Linagliptin treatment.** Linagliptin was formulated into the western diet at concentration of 83mg/kg. $\beta$ARKO$^{MIP}$ and control mice received either western diet without linagliptin or linagliptin diet at week 18. Metabolic tests were performed after 4 weeks of linagliptin treatment.

**Mouse islet steroid conversion assays.** Mouse islets were isolated from 10 wild-type male mouse and recovered overnight in complete medium: RPMI-1640 (Gibco) supplemented with 10% charcoal-stripped FBS and Pen/Strep (100 U/ml, 100 µg/ml). Approximately 250 islets were in each replicate and each condition was run in triplicate. Islets were treated with T (100nM; Sigma), the 5$\alpha$-R inhibitors finasteride (100nM; Sigma) and dutasteride (100nM; Sigma), or vehicle (ethanol and DMSO). Other control conditions included culture medium without FBS and complete medium with finasteride and dutasteride. Culture medium and islets were harvested for further analysis after a 24-hour incubation period.

**Steroid quantification by ultra-high-performance liquid-chromatography tandem mass spectrometry (UHPLC-MS/MS).** For the measurement of androgens, 500 µL of culture medium or external standard mix were combined with an internal standard mixture and extracted by liquid-liquid extraction with tert-butyl methyl ether (Acros Organics). Chromatographic separation and steroid quantification were performed using an ACQUITY ultra performance liquid chromatography system (Waters) coupled to a XEVO™ TQ-XS triple quadrupole mass spectrometer (Waters). Mass-to-charge transitions monitored in multiple reaction monitoring used for quantification are summarized in **Table S2**. Peak area ratios of analyte and internal standard, 1/x weighting and linear least square regression were used to produce the standard curves for quantification. Limits of quantifications were 0.24 nM for T and 0.24 nM for 5α-dihydrotestosterone (DHT).

**Statistical analysis.** Statistical analyses were performed with GraphPad Prism. Results are presented as mean ± SEM as specified in figures. When results showed a Gaussian distribution, statistical analyses were performed using the unpaired Student's *t* test or ANOVA (with Bonferroni post hoc test). A P value less than 0.05 was considered statistically significant. * $P<0.05$, ** $P<0.01$, ***$P < 0.001$, ****$P < 0.0001$.
